# Epstein-Barr virus noncoding RNA EBER1 promotes the expression of a ribosomal protein paralog to boost oxidative phosphorylation

**DOI:** 10.1101/2024.06.15.599158

**Authors:** Sita Paudel, Nara Lee

**Affiliations:** Department of Microbiology and Molecular Genetics University of Pittsburgh School of Medicine, Pittsburgh, PA 15219, USA

## Abstract

Epstein-Barr virus (EBV) is a highly successful pathogen that infects ∼95% of the adult population and is associated with diverse cancers and autoimmune diseases. The most abundant viral factor in latently infected cells is not a protein but a noncoding RNA called EBV-encoded RNA 1 (EBER1). Even though EBER1 is highly abundant and was discovered over forty years ago, the function of EBER1 has remained elusive. EBER1 interacts with the ribosomal protein L22, which normally suppresses the expression of its paralog L22-like 1 (L22L1). Here we show that when L22 binds EBER1, it cannot suppress L22L1, resulting in L22L1 being expressed and incorporated into ribosomes. We further show that L22L1-containing ribosomes preferentially translate mRNAs involved in the oxidative phosphorylation pathway. Moreover, upregulation of L22L1 is indispensable for growth transformation and immortalization of resting B cells upon EBV infection. Taken together, our results suggest that the function of EBER1 is to modulate host gene expression at the translational level, thus bypassing the need for dysregulating host gene transcription.

## Introduction

Epstein-Barr virus (EBV) was the first human cancer–causing virus discovered, and we now know that EBV is associated with several cancers (i.e., Burkitt, non-Hodgkin, and Hodgkin lymphomas; nasopharyngeal carcinoma; and gastric cancers) ^1–4^, accounting for ∼2 of every 100 cancer deaths ^5^. EBV infection may also drive serious and prominent immune diseases, like multiple sclerosis and Long COVID ^6,7^. Despite long-standing attention to the cellular events underlying EBV infection, we still lack treatments or vaccines for EBV-related diseases.

EBV expresses many noncoding RNAs ^8^. Of these, EBV-encoded RNA 1 (EBER1) is the most abundant viral factor during latency (about a million molecules per cell). Despite the field’s keen interest in EBER1 and the four decades since its discovery ^9^, we have yet to understand why EBV expresses EBER1. Efforts were made to examine how depleting or deleting EBER1 affects the virus or the host. But all earlier EBER1-depletion experiments failed because the RNA is impervious to antisense oligonucleotide–mediated silencing ^10^. And although EBER1-deletion strains were generated and used in *in vitro* infection studies (i.e., growth-transformation assays), deletion of EBER1 manifested no overt phenotype ^11,12^. This starkly contrasts with the fact that EBER1 is preserved and highly conserved in every clinical isolate ^13,14^, indicating that EBV undoubtedly needs EBER1 during natural infection. To reconcile this discrepancy, it was proposed that other EBV factors may compensate for EBER1 deletion *in vitro*, allowing deletion strains to immortalize B cells in infection assays ^15^.

Even though we have poorly understood EBER1’s molecular mode of action, its expression has clear consequences for host cells. EBER1 can drive oncogenesis, as its expression is sufficient to form colonies on soft agar and tumors in animals ^16^. Cells expressing exogenous EBER1 also have higher mitochondrial activity and are more proliferative than cells expressing no EBER1 ^17^. Astonishingly, EBER1 is functionally equivalent to TMER4, a noncoding RNA of similar length and secondary structure expressed by the EBV-related murine gammaherpesvirus 68. EBER1 can rescue TMER4 deletion to promote the egress of infected B cells from lymph nodes into peripheral circulation ^18^.

Known EBER1-interacting proteins are the RNA chaperone La, the RNA stability factor AUF1, the RNA methyltransferase NSUN2, and—most important for this study—the ribosomal protein L22 (**Supplementary Fig. 1a**) ^9,10,19,20^. L22 is a core protein in the large ribosomal subunit. The position of L22 in the large ribosomal subunit (**Supplementary Fig. 1b**) does not immediately make clear why L22 binding EBER1 would be significant; L22 is near neither the small subunit– large subunit interface nor the protein exit tunnel. The most striking L22 phenotype regarding EBER1 is that L22 relocalizes from nucleoli to the nucleoplasm when EBER1 is expressed ^11,20^, yet the significance of EBER1 binding L22 has remained unclear. A paralog of L22 called L22-like1 (L22L1) is encoded in the genome, and L22 has been reported to suppress the expression of L22L1 ^21^. The paralogs have distinct functions ^22^, particularly during hematopoiesis. This is intriguing because inclusion of sub-stoichiometric ribosomal proteins or incorporation of ribosomal protein paralogs produce so-called specialized ribosomes ^23^, which may preferentially translate subsets of mRNAs.

In this study, we found that EBER1 inhibits L22 and thus upregulates L22L1. When activated, L22L1 is incorporated into specialized ribosomes that preferentially translate mRNAs involved in the oxidative phosphorylation pathway (OXPHOS). Finally, we found that L22L1 expression is physiologically relevant during EBV infection and critical for EBV to immortalize B cells. Overall, these results suggest that EBER1 controls gene expression not by affecting transcription, but by producing specialized ribosomes to impact translation of pre-existing cellular mRNAs.

## Materials and Methods

### Purification of ribosomes by ultracentrifugation over a sucrose cushion

Ribosomes were purified as previously described with minor modifications ^23^. BJAB-B1 cells (2 x 10^7^ cells) were washed with PBS and lysed in 300 μl Sucrose Lysis Buffer (10 mM Tris pH 8.0; 0.32 M sucrose; 3 mM CaCl_2_; 2 mM magnesium acetate; 0.1 mM EDTA; 0.1% NP-40; 2 mM DTT; 1X protease inhibitor cocktail; 1 mM PMSF) and incubated on ice for 10 min. Cell lysate was spun for 10 min at 750 g in a tabletop centrifuge at 4°C to pellet cell nuclei. Supernatant was transferred to a fresh tube and spun for 10 min at 12,500 g at 4°C to pellet mitochondria. KCl was added to the post-mitochondrial cytoplasmic fraction to a final concentration of 0.5 M. Adjusted lysate was overlaid on 2 ml of sucrose cushion (50 mM Tris pH 7.4; 1 M sucrose; 5 mM MgCl_2_; 0.5 KCl) in a SW50Ti tube and ultracentrifuged for 2 h at 46,000 rpm at 4°C. After the run, supernatant was discarded, ribosomal pellet was carefully rinsed twice with ice-cold 200 μl H_2_O, and resuspended in 200 μl of Ribosome Resuspension Buffer (50 mM Tris pH 7.4; 5 mM MgCl_2_; 25 mM KCl).

### CRISPR-Cas9 mediated genome editing of BJAB-B1 cells

BJAB-B1 cells (2 x 10^6^ cells) were nucleofected using Lonza’s 4D-Nucleofector with SF solution and DS-120 program. Two plasmids were introduced, one of which expressed SpCas9 (0.5 μg of plasmid pX165 obtained from Addgene); the other plasmid (2.0 μg, in a pBluescript backbone) expressing an sgRNA targeting the 3′ region of the *RPL22L1* gene adjacent to the stop codon, and two homology-directed repair regions (one including an Avi-tag insertion) flanking a puromycin cassette. Two days after nucleofection, cells were cultured in the presence of 2 μg/ml puromycin, then diluted and seeded in a 96-well plate. Each well contained 5-10 cells, as BJAB-B1 cells will not grow from a single-cell culture. Individual cell clones were examined for proper gene editing by PCR analysis of genomic DNA, Sanger sequencing of the PCR amplicon, and by immunoblotting.

### Ribosome profiling and data analysis

Ribosome profiling was performed as previously described ^24^ with minor adjustments. CRISPR-edited BJAB-B1 cells expressing Avi-tagged L22L1 and BirA ligase were cultured to a density of ∼10^6^ cells/ml. Harringtonine was added at a final concentration of 2 µg/ml to the culture medium (40 ml, ∼4 x 10^7^ cells), and cells were treated for 5 min at 37°C. Cells were spun down at 3,000 rpm for 1 min at 4°C and medium was discarded. The cell pellet was washed with ice-cold PBS, then resuspended in 200 µl of Sucrose Lysis Buffer followed by a 10-min incubation on ice. Lysate was spun in a tabletop centrifuge at 3,000 rpm for 5 min at 4°C, and 600 µl of Lysis Dilution Buffer (20 mM Tris pH 7.4; 150 mM NaCl; 5 mM MgCl_2_; 1 mM DTT) was added to the supernatant. Lysate was centrifuged in a tabletop centrifuge for 3 min at full speed to pellet debris, nuclei, and mitochondria. DNase I (5 µl of 2U/µl) and RNase I (0.5 µl of 10U/µl) was added to the lysate and incubated for 45 min at RT with constant agitation in a thermoshaker. SUPERase•In (2 μl of 20 U/ µl) was added, and the lysate was cooled on ice. Lysate was overlaid on 2.5 ml of sucrose cushion in a Beckman SW50Ti rotor tube and then ultracentrifuged for 2 h at 47,000 rpm at 4°C. Supernatant was discarded, 1 ml of TRIzol was added to the ribosomal pellet to isolate RNA. RNA was resolved in a 15% denaturing polyacrylamide gel, and RNA fragments of 24-34 nts were gel-extracted. Size-selected RNA fragments were treated with polynucleotide kinase to remove 2′-3′ cyclic phosphate and add 5′ phosphate by mixing 14 μl of RNA, 2 μl of 10x PNK Buffer, 2 μl of 10 mM ATP, and 2 μl of T4 PNK in a 20 μl-reaction and incubating for 30 min at 37°C. RNA was then phenol-chloroform-extracted, followed by rRNA depletion using a mix of six antisense oligonucleotides (ASOs) complementary to rRNA (see **Supplementary Table 1**). For this, RNA was resuspended in 7 µl H_2_O, rRNA ASOs were prepared at 10 µM each; 2 µl of ASO mix and 1 µl of 20x SSC were added. Solution was heated for 90 sec at 100°C, and rRNA and ASOs were annealed by decreasing the temperature at - 0.1°C/sec to 37°C, followed by an incubation at 37°C for 15 min. In the meantime, 25 µl of MyOne Streptavidin C1 beads were washed three times with 1X Wash/Bind Buffer (5 mM Tris pH 7.4; 1 M NaCl; 1 mM EDTA; 0.2% Triton X-100). 10 µl of 2X Bind/Wash Buffer were added to beads, pre-warmed to 37°C, and RNA solution was added for 15 min at 37°C with constant mixing at 1,000 rpm in a thermoshaker. Beads were precipitated and 17.5 µl of eluate was recovered and ethanol-precipitated. RNA was resuspended in 7.5 μl H_2_O, concentration was measured by Qubit, and an Illumina-compatible library was generated using the NEBNext Small RNA Library Prep Set. To prepare Ribo-seq libraries from L22L1-containing ribosomes specifically, L22L1-ribosomes were pulled down with 25 μl of MyOne T1 streptavidin beads after DNase I/RNase I digestion, and RNA was isolated for size-selection by PAGE using phenol-chloroform extraction. Deep sequencing data generated in this study were deposited in the Sequence Read Archive under BioProject ID PRJNA1093137. Ribo-seq data was analyzed using Qiagen’s CLC Genomics Workbench. Please note that Ribo-seq data were analyzed as RNA-seq data without normalizing for mRNA expression levels because total ribosomes and Avi-tagged L22L1-containing ribosomes were isolated from the same starting cell line and are thus internally controlled. Differential expression data was then analyzed with Qiagen’s Ingenuity Pathway Analysis to identify affected gene categories.

### Polysome fractionation by sucrose gradient ultracentrifugation and qRT-PCR analysis

Polysome fractionation was performed as previously described ^25^. In brief, BJAB-B1 cells were treated with 100 µg/ml cycloheximide for 10 min in a CO_2_-incubator at 37°C. Cells were washed with ice-cold PBS and lysed with 400 µl of Lysis Buffer (10 mM HEPES pH 7.4; 150 mM KCl; 10 mM MgCl_2_; 1 mM DTT; 2% NP-40; 1X protease inhibitor cocktail; RNase inhibitor 6 U/ml). Cell lysate was loaded on a 15-45% sucrose gradient in SW41-rotor tubes and ultracentrifuged at 40,000 rpm for 2 h at 4°C. The gradient was then fractionated using BioComp Instrument’s Piston Gradient Fractionator following manufacturer’s instructions. RNA from the indicated polysome fractions was isolated by phenol-chloroform extraction. Isolated RNA was converted into cDNA for subsequent qRT-PCR analysis. Primer sequences are listed in **Supplementary Table 1**.

### Measurement of ATP levels

BJAB-B1 cells and derivative knockdown cell lines for EBER1 or L22L1 were used to measure intracellular ATP levels using PerkinElmer’s ATPlite Luminescence ATP Detection Assay System following manufacturer’s instructions.

### EBV-induced growth transformation of primary B cells coupled with CRISPRi

Growth transformation of resting B cells were performed as previously described ^26^. To prepare EBV stocks, marmoset B95-8 cells were grown in RPMI 1640 medium to a concentration of 10^6^ cells/ml and then stimulated with 20 ng/ml TPA (tetradecanoylphorbol acetate) for 1 h. Cells were washed three times with medium, resuspended in the original volume, and incubated for 96 h before harvesting the supernatant by centrifuging the culture at 600 g for 10 min at 4°C. The supernatant was filtered through a 0.45-micron filter, aliquoted, and stored at -80°C.

PBMCs were obtained from Lonza, and 5 x 10^6^ cells were thawed and resuspended at a concentration of 2 x 10^6^ cells/ml in RPMI 1640 medium for each *in vitro* infection assay. FK506 (final concentration of 20 nM) and polybrene (final concentration of 5 µg/ml) were added before CRISPRi lentiviruses at a MOI of 5 were added to the medium for 1 h in a CO_2_-incubator at 37°C. Then, EBV infectious particle-containing stock was thawed and added to the medium at 1/10 of the total volume. Flask was swirled gently and placed in a CO_2_-incubator at 37°C for several weeks and examined as indicated.

## Results

### EBER1 prevents L22 from suppressing L22L1

Because EBER1 is impervious to hybridization-based knockdown approaches, we tested whether we could use CRISPRi to deplete EBER1. We designed single-guide (sg)RNAs targeting the EBER1 locus and co-expressed them with dCas9-KRAB in EBV-infected BJAB-B1 cells. With this CRISPRi approach, we reduced EBER1 expression to less than 10% of its normal levels (**Fig. 1a**). We then analyzed EBER1-depleted cells by RNA-seq to determine whether EBER1 transcriptionally affects any specific pathway. Even though 181 and 183 genes were more than twofold down and upregulated (p<0.05), respectively (**Supplementary Table 2**), we found no particular category or pathway affected by using gene ontology analysis. However, one downregulated gene was *RPL22L1*, which encodes the L22 paralog L22L1 (**Fig. 1b**, **Supplementary Fig. 1c**). By qRT-PCR, we verified that EBER1 depletion lowered levels of *RPL22L1* mRNA (**Fig. 1c**). Because no paralog-specific antibody was available, we purified ribosomes from control or EBER1-depleted cells and performed mass spectrometry analysis to determine whether protein levels are similarly affected. Overall ribosome composition in control cells did not differ from that in EBER1-depleted cells except that EBER1-depleted cells lacked L22L1 (**Supplementary Table 3**). To verify this result, we produced our own anti-L22L1 antibody and confirmed by immunoblotting that ribosomes from EBER1-depleted cells had no L22L1 (**Fig. 1d**). By applying parallel reaction monitoring with labeled internal standard peptides (a mass-spec method for absolute quantification of proteins), the ratio of L22 to L22L1 in EBER1-expressing cells was found to be 31:1. This indicated that the majority of ribosomes in EBER1-expressing cells contain L22, but that a subset has incorporated L22L1. Regarding the regulation of L22L1, it was reported that the canonical ribosomal protein L22 inhibits L22L1 expression ^21^. We verified this previous finding by depleting L22 and observing increased L22L1 expression when L22 is no longer present to enforce its suppressive action (**Supplementary Fig. 1d**). Taken together, these data indicate that when EBER1 binds L22, it prevents L22 from suppressing L22L1, upregulates L22L1, and lets L22L1 be incorporated into a subset of ribosomes.

**Figure 1.**
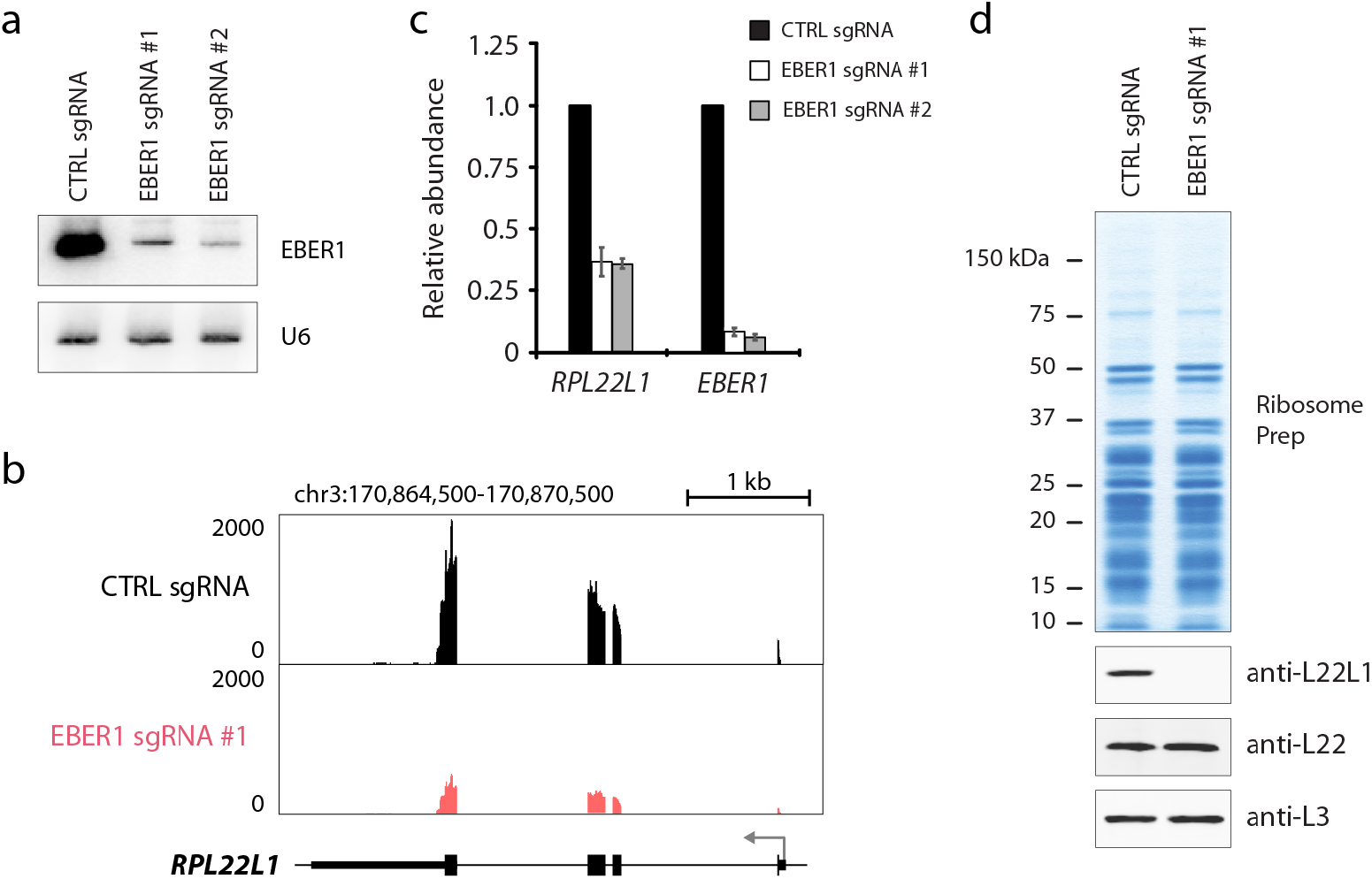
EBER1 expression results in the upregulation of L22L1 and its incorporation into ribosomes. (**a**) Knockdown of EBER1 by using CRISPRi in EBV-positive BJAB-B1 cells. Two sgRNAs were used to target dCas9-KRAB to the EBER1 locus on the EBV genome. Both sgRNAs deplete EBER1 to less than 10% of wildtype levels. (**b+c**) RNA-seq result for *RPL22L1* after EBER1 knockdown in BJAB-B1 cells (b). Please see **Supplementary Table 1** for further information. *RPL22L1* is downregulated upon EBER1 depletion, which was confirmed by qRT-PCR analysis (c). (**d**) EBER1-expressing cells express L22L1 and have L22L1-containing ribosomes. Ribosomes were isolated from control and EBER1-knockdown cells and subjected to L22L1 immunoblot analysis. The cell line expressing sgRNA #1 is shown as a representative. Antibodies against L22 and L3 were used as controls. EBER1 depletion abrogates L22L1 expression and ribosome incorporation. These ribosome preparations were analyzed by mass spectrometry (**Supplementary Table 2**), which showed that the main difference in ribosome composition upon EBER1 depletion is with L22L1 only.

### L22L1-containing ribosomes preferentially translate OXPHOS mRNAs

To examine whether mRNAs translated by L22L1-containing ribosomes differ from those translated by canonical ribosomes (containing L22), we isolated L22L1-containing ribosomes from EBV-positive cells and compared their translatome with that of the total pool of ribosomes. Because a C-terminal HA-tag is suitable for purifying L22-containing ribosomes ^27^ and because this region is highly conserved between L22 and L22L1 (**Supplementary Fig. 1c**), we reasoned that we could purify L22L1-containing ribosomes by expressing C-terminally HA-tagged L22L1. We CRISPR-engineered EBV-positive BJAB-B1 cells to express HA-tagged L22L1 from the endogenous gene locus. However, while HA-tagged L22 is functional in precipitating whole ribosomes, the addition of an HA-tag to L22L1 did not facilitate selection of whole ribosomes (**Supplementary Fig. 2b**), and neither did the addition of a C-terminal FLAG-tag to L22L1 (data not shown). We next considered using an Avi-tag because previous reports applied an Avi-tag insertion to ribosome-associated factors, which become biotinylated by the co-expressed BirA ligase, and these ribosomes were successfully isolated with streptavidin beads for subsequent Ribo-seq analysis ^28,29^. We CRISPR-edited an Avi-tag at the C-terminus of L22L1 and generated BJAB-B1 cell lines that stably co-express BirA ligase (**Fig. 2a-c**). Using streptavidin beads, we could precipitate whole L22L1-containing ribosomes (**Supplementary Fig. 2a**), as indicated by immunoblotting for another ribosomal protein (L3; **Fig. 2d**) and by mass-spec analysis of streptavidin-bead precipitate (**Supplementary Table 4**). Importantly, selecting with streptavidin beads specifically enriched for L22L1-containing ribosomes, while largely discarding L22-containing ones (**Fig. 2d**). We then performed ribosome profiling with either the total pool of ribosomes or specifically with L22L1-containing ribosomes. Principle component analysis of these data showed that L22L1-containing ribosomes are functionally distinct from the total pool of ribosomes, as the replicates grouped into separate clusters (**Fig. 2e**). Ingenuity Pathway Analysis of differentially translated mRNAs showed that the L22L1-containing ribosomes most affected the oxidative phosphorylation (OXPHOS) pathway and preferentially translated OXPHOS mRNAs compared to the total pool of ribosomes (**Fig. 2f**).

**Figure 2.**
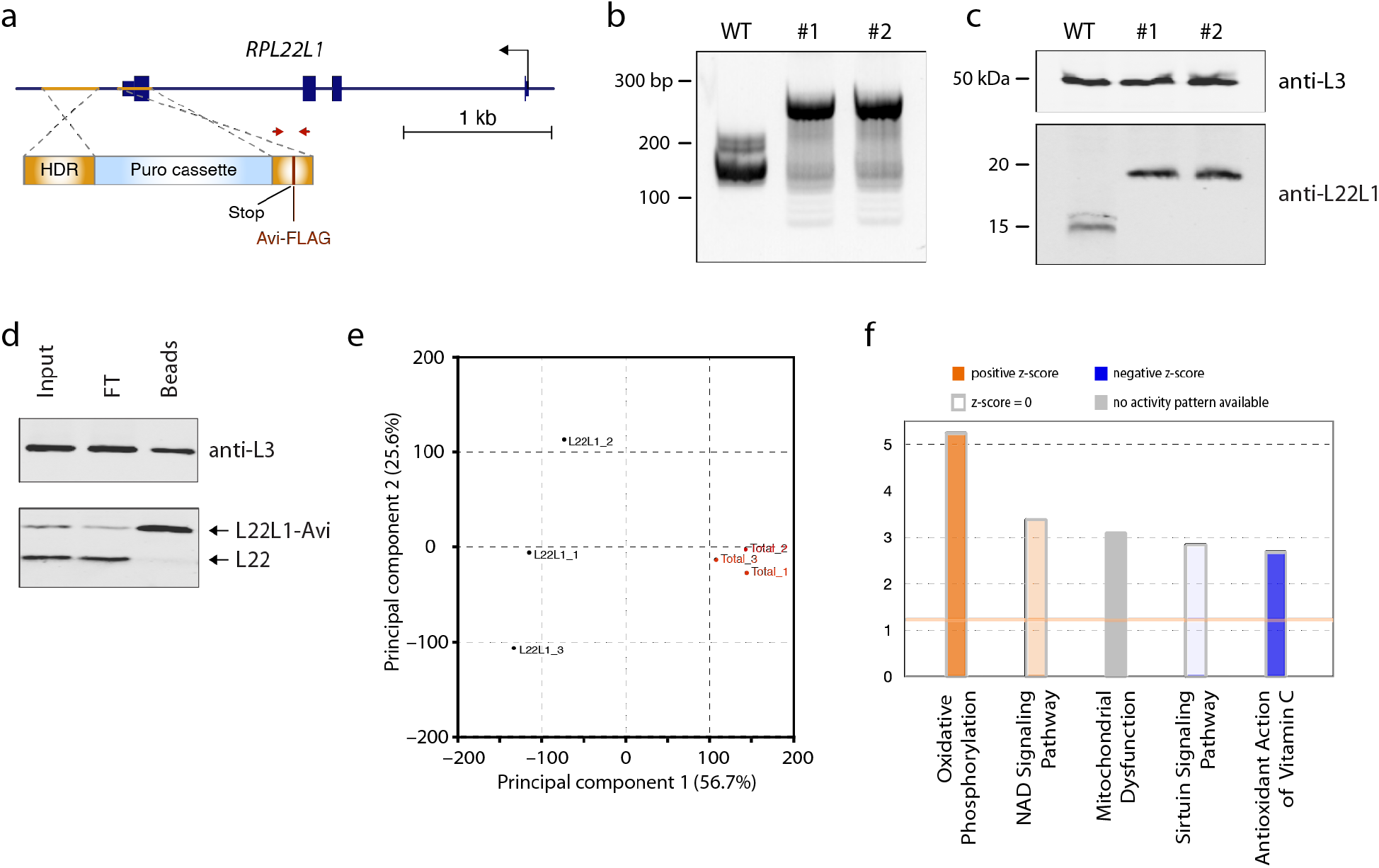
L22L1-containing ribosomes preferentially translate mRNAs involved in the oxidative phosphorylation pathway. (**a**) CRISPR-mediated homology-directed repair approach for inserting a C-terminal Avi-tag into the endogenous *RPL22L1* locus in BJAB-B1 cells. A puromycin cassette was included to facilitate with the selection of edited cell clones. A FLAG-tag was also included for downstream detection. Arrows indicate regions to which primers anneal for PCR amplification from genomic DNA. (**b**) PCR amplification of the edited region from control and two edited cell lines. The shift in band size corresponds to sequences encoding the Avi and FLAG-tag inserted into the 3′ region of *RPL22L1*. The PCR amplicons were analyzed by Sanger sequencing to confirm correct editing. (**c**) Insertion of Avi-FLAG tag was confirmed in two CRISPR-edited cell lines by immunoblotting, indicated by the shift in band size. Anti-L3 antibody was used as a control. (**d**) Purification of L22L1-containing ribosomes by using streptavidin beads. 10% of input and 10% of flow-through (FT), and 100% of the precipitated material were loaded on the gel. L22L1-containing ribosomes were highly enriched, while canonical L22-containing ribosomes were not. Please note that a substantial amount of L3 is found in the FT, which is consistent with the fact that most ribosomes contain L22 and not L22L1 (31:1 ratio). (**e**) Principal component analysis of Ribo-seq data comparing the translatome of total ribosomes with that of L22L1-containing ones. The sample sets cluster in distinct groups, indicating distinct cohorts of translated mRNAs. (**f**) Ingenuity Pathway Analysis indicates that OXPHOS mRNAs are preferentially translated by L22L1-containing ribosomes compared to canonical L22-containing ones.

To confirm our Ribo-seq results, we measured the levels of OXPHOS mRNAs on translating ribosomes from control cells and from EBER1-depleted EBV-positive BJAB-B1 cells. We first separated ribosome fractions by polysome fractionation. Overall distributions of subunits, whole ribosomes, and polysomes did not markedly change upon EBER1 depletion (**Fig. 3a**). The distributions of L22 and L22L1 in control cells were also similar (**Fig. 3b**). We then isolated total RNA from polysome fractions and measured the abundance of candidate OXPHOS mRNAs by qRT-PCR analysis. While total mRNA levels were comparable in control and EBER1-knockdown cells, OXPHOS mRNAs were significantly more abundant (more than twofold) in polysome fractions from control cells than they were upon EBER1 knockdown (**Fig 3c**), suggesting that EBER1 regulates translation of OXPHOS mRNAs. To further test whether EBER1 impacts OXPHOS, we measured the intracellular ATP levels after EBER1 or L22L1 knockdown. Consistent with EBER1 regulating OXPHOS, depletion of either EBER1 or L22L1 resulted in decreased ATP levels in BJAB-B1 cells (**Fig. 3d**). Taken together, mitochondrial function measurements support our Ribo-seq result that L22L1 upregulates the OXPHOS pathway.

**Figure 3.**
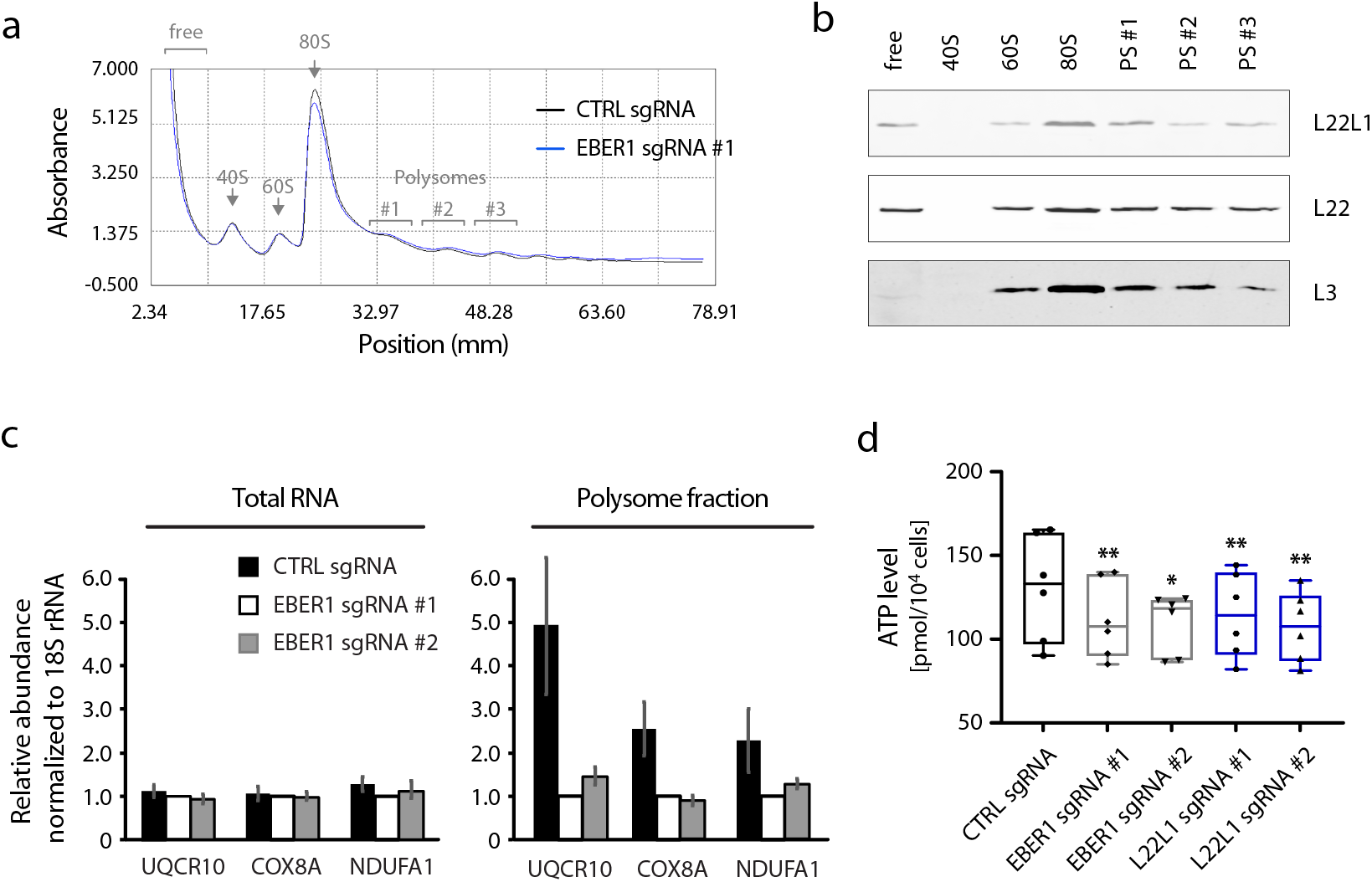
OXPHOS mRNAs are enriched on translating ribosomes of EBER1-expressing cells. (**a**) Polysome fractionation of control and EBER1-knockdown BJAB-B1 cell lysates by ultracentrifugation over sucrose gradients. Distinct ribosomal subunits (free, 40S, 60S, 80S, and three different polysome fractions) are indicated. EBER1 knockdown did not change overall ribosomal subunit distributions. (**b**) L22 and L22L1 show a comparable distribution in ribosome/polysome fractions. Individual fractions, as indicated in (a), were subjected to immunoblotting for L22L1, L22, and L3. Consistent with earlier reports, L22 and L22L1 were found in ribosome-free fractions, but not L3. (**c**) qRT-PCR analysis of candidate OXPHOS mRNAs in total or polysome-associated RNA from control or EBER1-depleted BJAB-B1 cells. RNA was isolated from intact cells (total RNA) or from polysome fractions #1–3 , as indicated in (a). qRT-PCR was performed for *UQCR10*, *COX8A*, and *NDUFA1*. These mRNAs were enriched in polysomes of EBER1-expressing cells compared to EBER1-depleted cells. RNA expression levels for these mRNAs did not change upon EBER1 knockdown, indicating a translational upregulation. (**d**) Decreased ATP levels in BJAB-B1 cells depleted for either EBER1 or L22L1. Consistent with preferential translation of OXPHOS mRNAs, either EBER1 knockdown or L22L1 knockdown by CRISPRi with two sgRNAs lowered intracellular ATP levels. **, p<0.01; *, p=0.02 by paired t-test.

### EBV needs L22L1 to transform B cells

To examine whether L22L1 upregulation is critical for EBV to infect cells, we performed growth-transformation assays with concomitant knockdown of L22L1. In this approach, resting B cells in peripheral blood mononuclear cells (PBMCs), which do not grow in culture, are infected with EBV, upon which they become immortalized and start proliferating. We infected PBMCs with both EBV infectious particles and lentiviral L22L1-knockdown constructs (**Fig. 4a, b**). These single-vector lentiviral knockdown constructs expressed sgRNAs targeting the *RPL22L1* promoter, dCas9-KRAB, and a puromycin cassette. Three weeks after infection, control knockdown cells formed colonies and proliferated in culture, but L22L1-knockdown PBMCs did not (**Fig. 4c**). To confirm that outgrowing cells carried the lentiviral knockdown construct, we expanded them under puromycin selection and immunoblotted for HA-tagged dCas9-KRAB (**Fig. 4d**). We also confirmed that these cells were infected with EBV by showing that they expressed EBNA1 and EBER1 (**Fig. 4d**). Taken together, these results indicate that L22L1 upregulation is critical for EBV to immortalize B cells.

**Figure 4.**
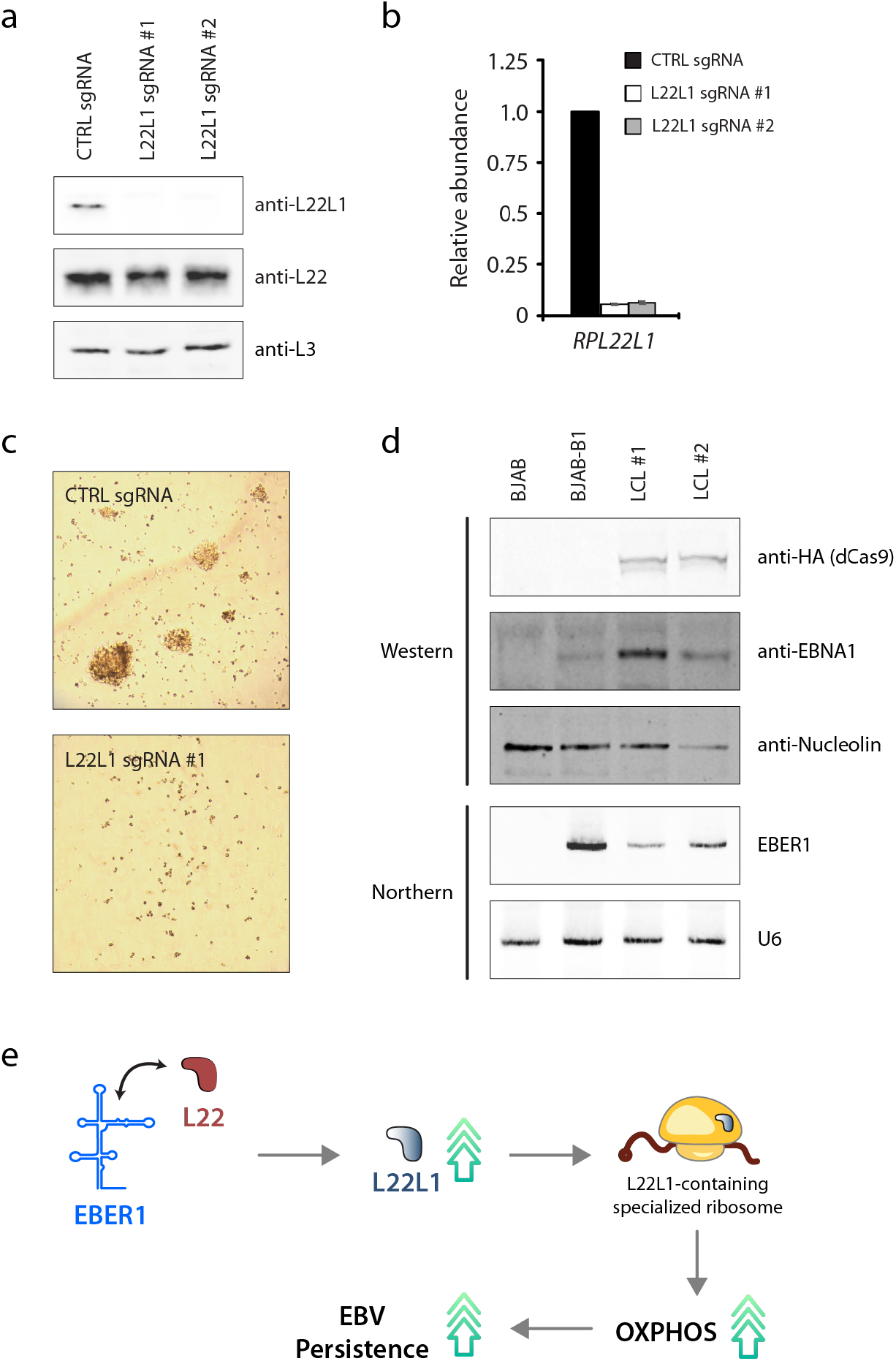
L22L1 is essential for EBV-induced growth transformation of primary B cells. (**a**) L22L1 is efficiently knocked down by using two lentiviral constructs expressing distinct sgRNAs and dCas9-KRAB. The same lentiviral vector also expressed a puromycin resistance gene. (**b**) *RPL22L1* mRNA levels were measured by qRT-PCR analysis after knockdown with the same sgRNAs as shown in (a), confirming efficient depletion of *RPL22L1*. (**c**) Colony formation of EBV-infected PBMCs. Three weeks after EBV infection and transduction with L22L1-knockdown lentivirus, only control-knockdown cells formed colonies and proliferated in culture. PBMCs transduced with sgRNA #1 only are shown; a second knockdown cell line similarly did not show cell outgrowth. (**d**) Two independently generated LCLs were examined for the expression of dCas9-KRAB (by immunoblotting with an anti-HA-tag antibody), and for expression of EBNA1 and EBER1 to confirm EBV infection. Nucleolin and U6 were as loading controls. Cell lysates or RNA from BJAB or BJAB-B1 cells were references. (**e**) Model for EBER1–L22 interaction increasing OXPHOS activity and EBV persistence by inducing L22L1-containing ribosomes.

## Discussion

A long-standing mystery in the EBV and noncoding-RNA fields is the function of EBER1 RNA. In this study, we show that EBER1 affects host gene expression by regulating translation (**Fig. 4e**). By binding L22, EBER1 prevents L22 from suppressing its paralog, L22L1. L22L1 is then expressed and incorporated into specialized ribosomes, which preferentially translate OXPHOS mRNAs. Finally, we show that upregulated L22L1 in infected cells is essential for EBV to transform B cells. Thus, EBER1 regulates gene expression at the level of translation, changing translation levels of existing cellular mRNAs, which differs from how many other prominent noncoding RNAs regulate gene expression, i.e., at the level of transcription by binding transcription factors ^30^.

Our study is not the first to link EBV infection to OXPHOS upregulation. A previous study has shown that OXPHOS is upregulated in resting B cells, as they become immortalized by EBV infection ^31^. Based on our results, EBER1 is the factor that executes this important regulatory step. EBER1 is conserved in every clinical isolate and thus probably critical for the EBV life cycle *in vivo*. That EBV does not need EBER1 to immortalize B cells *in vitro* is therefore puzzling. EBER1 may be essential *in vivo,* but not *in vitro*, if redundant EBV factors can induce L22L1 expression. For example, this redundant pathway would not be activated during natural infection, but can be activated during artificial infection of cultured cells. Still, our results suggest that L22L1, which is downstream of EBER1, is both essential for growth transformation and a rate-limiting factor in this process. That L22L1 is critical for EBV infection is supported by the fact that *RPL22L1* is expressed in EBV-transformed lymphocytes more highly than in any other tissue tested (**Supplementary Fig. 3**).

While our results indicate that L22L1-containing ribosomes increase OXPHOS activity, which has been shown to be necessary for the metabolic reprogramming during LCL generation ^31^, key questions remain. First, it is unclear how these specialized ribosomes target OXPHOS mRNAs for translation. The small ribosomal subunit with the help of auxiliary factors handles translation initiation and mRNA scanning; the large subunit, of which L22L1 is part, is assembled only at later stages. Thus, L22L1 may not affect how ribosomes assemble onto mRNAs, but may instead increase the efficiency of ribosome translation or the stability of translating ribosomes, which is what we may be observing in our Ribo-seq data. Another outstanding question is the mechanism by which L22 inhibits L22L1. A simple model as previously proposed in which L22 binds and degrades *RPL22L1* mRNA ^21^ is unlikely because we found no evidence of L22 binding *RPL22L1* in our L22-CLIP data and because L22 has no RNase activity. Mere binding to a transcript would not explain degradation of that transcript. Indeed, L22 binds both EBER1 and 28S ribosomal RNA, but degrades neither.

Overall, these results show how EBER1 regulates translation. Although this is an elegant mechanism to affect host gene expression, EBER1 may have yet other molecular functions and regulate other pathways under certain circumstances during the complex EBV life cycle. We found that EBER1 robustly binds another novel protein (unpublished data), and additional studies are needed to fully dissect the molecular mechanisms of this intriguing viral ncRNA.

## Acknowledgements

We thank Dr. Joan Steitz (Yale University) for reagents and discussion; Dr. George Miller (Yale University) for reagents; Dr. Sumit Borah for editorial assistance. This work was supported by the National Institutes of Health [grant number R21AI151073 and R21AI163246] to NL.

## Author Contributions

S.P. performed experiments and analyzed data; N.L. designed the study, performed experiments, analyzed data, and wrote the manuscript.

## Supplementary Table

**Supplementary Table 1.** List of reagents used in this study.

**Supplementary Table 2.** RNA-seq analysis of control and EBER1-knockdown BJAB-B1 cells.

**Supplementary Table 3.** Mass-spec analysis of ribosomes isolated from control or EBER1-knockdown BJAB-B1 cells. Ribosome preparations from control cells differ from those from EBER1-knockdown cells by having L22L1.

**Supplementary Table 4.** Mass-spec analysis of L22L1-containing ribosomes selected with streptavidin beads. Streptavidin purification of Avi-tagged L22L1 precipitates whole ribosomes, as shown by the detection of proteins from both small and large ribosomal subunits.

## Supplementary Figure Legends

**Supplementary Figure 1.**
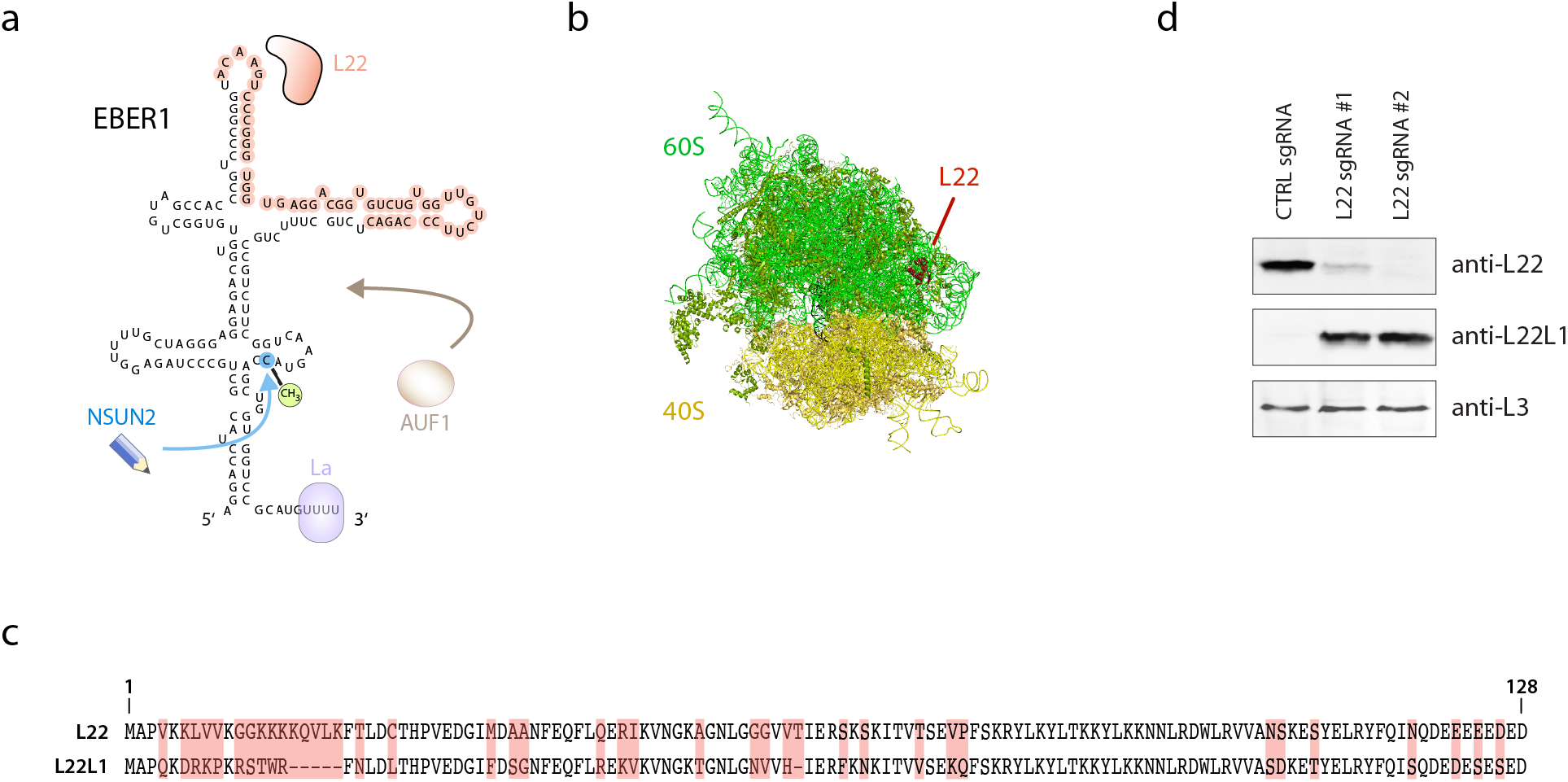
EBER1-interaction with L22 upregulates L22L1. (**a**) Secondary structure of EBER1 and its known interacting proteins are shown. The binding site of AUF1 is unknown. The binding site of L22 is shown in red. (**b**) The localization of L22 within the ribosome is indicated. The 60S subunit is colored in green, the 40S subunit in yellow. (**c**) Primary sequences of L22 and L22L1 are shown. Divergent amino acids are highlighted in red. The N-terminal regions show the highest degree of sequence variation. (**d**) Depletion of L22 upregulates L22L1. L22 was knocked down by using CRISPRi with two distinct sgRNAs targeting the *RPL22* promoter. As previously reported, depletion of L22 upregulated L22L1, confirming that L22 suppresses L22L1.

**Supplementary Figure 2.**
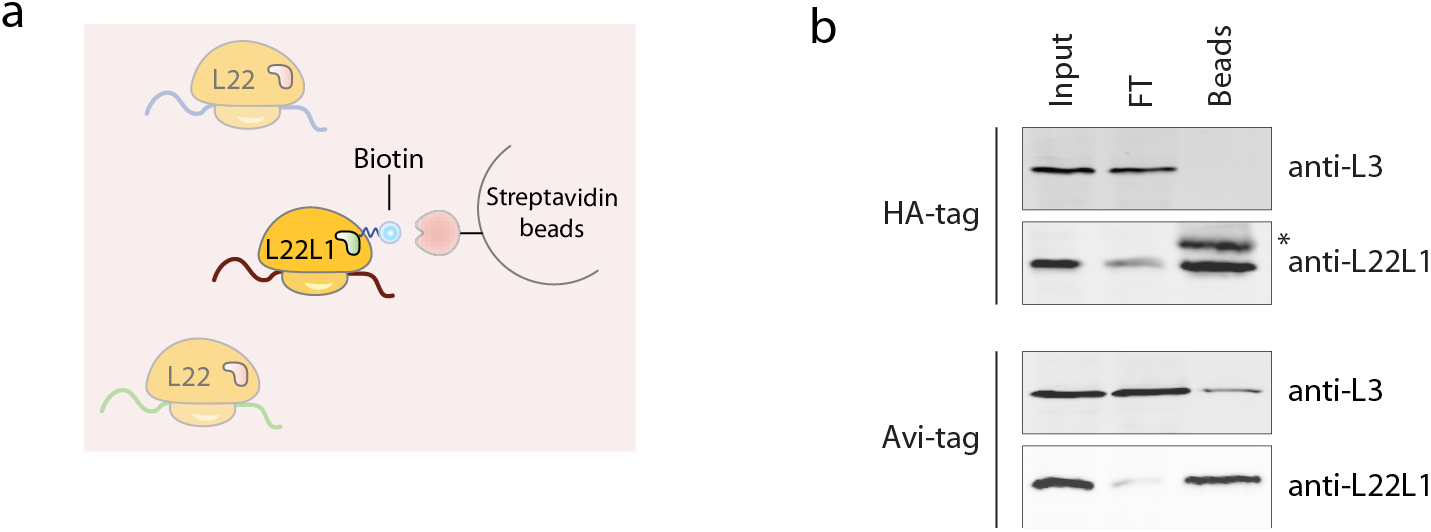
Isolating L22L1-containing ribosomes by using Avi-tagged L22L1. (**a**) Strategy for isolating L22L1-containing ribosomes from total pools of ribosomes. An Avi-tag is inserted into the C-terminus of L22L1 by CRISPR-mediated gene editing. BirA ligase is stably expressed in edited cells by lentiviral transduction, resulting in the biotinylation of L22L1 and subsequent purification by streptavidin beads. (**b**) Unlike earlier reports for L22, inserting an HA-tag at the C-terminus of L22L1 does not allow purification of whole ribosomes, but only of unincorporated L22L1 protein. This is shown by the absence of L3, another ribosomal protein of the large subunit, in the immunoblot (top panel set). Asterisk indicates the light chain of the HA-antibody used for co-IP. On the other hand, a C-terminal Avi-tag in L22L1 allows purification of whole ribosomes, as shown by the L3 immunoblot and as confirmed by mass-spec analysis (please see **Supplementary Table 4).**

**Supplementary Figure 3.**
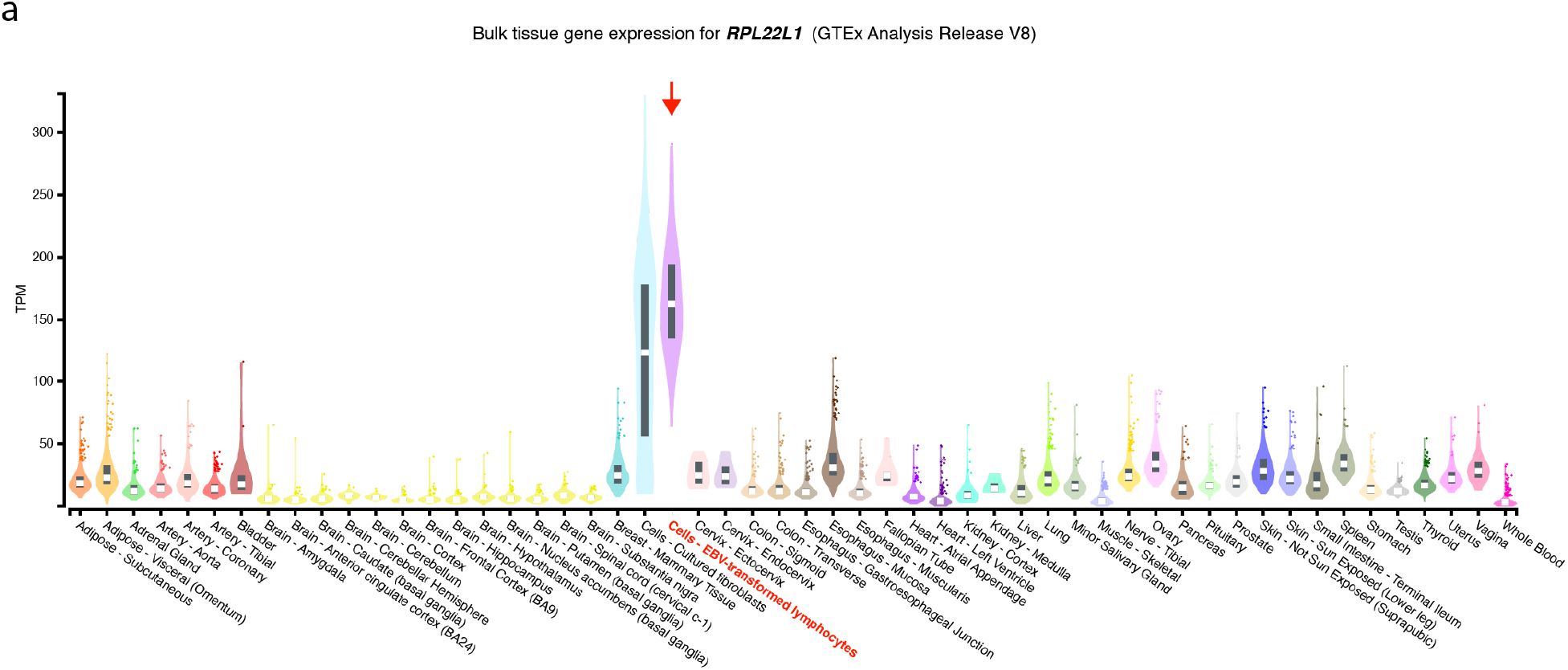
*RPL22L1* expression is highest in the category “Cells – EBV-transformed lymphocytes”. Data were taken from the GTex Analysis Release, version 8.

## Notes

### Competing Interest Statement

The authors have declared no competing interest.

## References

1 Young, L. S., Yap, L. F. & Murray, P. G. Epstein-Barr virus: more than 50 years old and still providing surprises. Nat Rev Cancer 16, 789–802, doi:10.1038/nrc.2016.92 (2016).

2 Hislop, A. D., Taylor, G. S., Sauce, D. & Rickinson, A. B. Cellular responses to viral infection in humans: lessons from Epstein-Barr virus. Annu Rev Immunol 25, 587–617, doi:10.1146/annurev.immunol.25.022106.141553 (2007).

3 Damania, B. Oncogenic gamma-herpesviruses: comparison of viral proteins involved in tumorigenesis. Nat Rev Microbiol 2, 656–668, doi:10.1038/nrmicro958 (2004).

4 Moore, P. S. & Chang, Y. Why do viruses cause cancer? Highlights of the first century of human tumour virology. Nat Rev Cancer 10, 878–889, doi:10.1038/nrc2961 (2010).

5 Zhong, L. et al. Urgency and necessity of Epstein-Barr virus prophylactic vaccines. NPJ Vaccines 7, 159, doi:10.1038/s41541-022-00587-6 (2022).

6 Bjornevik, K. et al. Longitudinal analysis reveals high prevalence of Epstein-Barr virus associated with multiple sclerosis. Science 375, 296–301, doi:10.1126/science.abj8222 (2022).

7 Su, Y. et al. Multiple early factors anticipate post-acute COVID-19 sequelae. Cell 185, 881–895 e820, doi:10.1016/j.cell.2022.01.014 (2022).

8 Lee, N. The many ways Epstein-Barr virus takes advantage of the RNA tool kit. RNA Biol 18, 759–766, doi:10.1080/15476286.2021.1875184 (2021).

9 Lerner, M. R., Andrews, N. C., Miller, G. & Steitz, J. A. Two small RNAs encoded by Epstein-Barr virus and complexed with protein are precipitated by antibodies from patients with systemic lupus erythematosus. Proc Natl Acad Sci U S A 78, 805–809 (1981).

10 Lee, N., Pimienta, G. & Steitz, J. A. AUF1/hnRNP D is a novel protein partner of the EBER1 noncoding RNA of Epstein-Barr virus. RNA 18, 2073–2082 (2012).

11 Gregorovic, G. et al. Cellular gene expression that correlates with EBER expression in Epstein-Barr Virus-infected lymphoblastoid cell lines. J Virol 85, 3535–3545 (2011).

12 Swaminathan, S., Tomkinson, B. & Kieff, E. Recombinant Epstein-Barr virus with small RNA (EBER) genes deleted transforms lymphocytes and replicates in vitro. Proc Natl Acad Sci U S A 88, 1546–1550 (1991).

13 Moss, W. N. & Steitz, J. A. Genome-wide analyses of Epstein-Barr virus reveal conserved RNA structures and a novel stable intronic sequence RNA. BMC Genomics 14, 543, doi:10.1186/1471-2164-14-543 (2013).

14 Xu, M. et al. Genome sequencing analysis identifies Epstein-Barr virus subtypes associated with high risk of nasopharyngeal carcinoma. Nat Genet 51, 1131–1136, doi:10.1038/s41588-019-0436-5 (2019).

15 Felton-Edkins, Z. A. et al. Epstein-Barr virus induces cellular transcription factors to allow active expression of EBER genes by RNA polymerase III. J Biol Chem 281, 33871–33880, doi:10.1074/jbc.M600468200 (2006).

16 Houmani, J. L., Davis, C. I. & Ruf, I. K. Growth-promoting properties of Epstein-Barr virus EBER-1 RNA correlate with ribosomal protein L22 binding. J Virol 83, 9844–9853 (2009).

17 Ahmed, W. et al. Epstein-Barr virus noncoding small RNA (EBER1) induces cell proliferation by up-regulating cellular mitochondrial activity and calcium influx. Virus Res 305, 198550, doi:10.1016/j.virusres.2021.198550 (2021).

18 Hoffman, B. A., Wang, Y., Feldman, E. R. & Tibbetts, S. A. Epstein-Barr virus EBER1 and murine gammaherpesvirus TMER4 share conserved in vivo function to promote B cell egress and dissemination. Proc Natl Acad Sci U S A 116, 25392–25394, doi:10.1073/pnas.1915752116 (2019).

19 Henry, B. A., Kanarek, J. P., Kotter, A., Helm, M. & Lee, N. 5-methylcytosine modification of an Epstein-Barr virus noncoding RNA decreases its stability. RNA 26, 1038–1048, doi:10.1261/rna.075275.120 (2020).

20 Toczyski, D. P., Matera, A. G., Ward, D. C. & Steitz, J. A. The Epstein-Barr virus (EBV) small RNA EBER1 binds and relocalizes ribosomal protein L22 in EBV-infected human B lymphocytes. Proc Natl Acad Sci U S A 91, 3463–3467 (1994).

21 O’Leary, M. N. et al. The ribosomal protein Rpl22 controls ribosome composition by directly repressing expression of its own paralog, Rpl22l1. PLoS Genet 9, e1003708, doi:10.1371/journal.pgen.1003708 (2013).

22 Zhang, Y. et al. Ribosomal Proteins Rpl22 and Rpl22l1 Control Morphogenesis by Regulating Pre-mRNA Splicing. Cell Rep 18, 545–556, doi:10.1016/j.celrep.2016.12.034 (2017).

23 Belin, S. et al. Purification of ribosomes from human cell lines. Curr Protoc Cell Biol Chapter 3, Unit 3 40, doi:10.1002/0471143030.cb0340s49 (2010).

24 Ingolia, N. T., Brar, G. A., Rouskin, S., McGeachy, A. M. & Weissman, J. S. The ribosome profiling strategy for monitoring translation in vivo by deep sequencing of ribosome-protected mRNA fragments. Nat Protoc 7, 1534–1550, doi:10.1038/nprot.2012.086 (2012).

25 He, S. L. & Green, R. Polysome analysis of mammalian cells. Methods Enzymol 530, 183–192, doi:10.1016/B978-0-12-420037-1.00010-5 (2013).

26 Hui-Yuen, J., McAllister, S., Koganti, S., Hill, E. & Bhaduri-McIntosh, S. Establishment of Epstein-Barr virus growth-transformed lymphoblastoid cell lines. J Vis Exp, doi:10.3791/3321 (2011).

27 Sanz, E. et al. Cell-type-specific isolation of ribosome-associated mRNA from complex tissues. Proc Natl Acad Sci U S A 106, 13939–13944, doi:10.1073/pnas.0907143106 (2009).

28 Shurtleff, M. J. et al. The ER membrane protein complex interacts cotranslationally to enable biogenesis of multipass membrane proteins. Elife 7, doi:10.7554/eLife.37018 (2018).

29 Jan, C. H., Williams, C. C. & Weissman, J. S. Principles of ER cotranslational translocation revealed by proximity-specific ribosome profiling. Science 346, 1257521, doi:10.1126/science.1257521 (2014).

30 Cech, T. R. & Steitz, J. A. The noncoding RNA revolution-trashing old rules to forge new ones. Cell 157, 77–94, doi:10.1016/j.cell.2014.03.008 (2014).

31 McFadden, K. et al. Metabolic stress is a barrier to Epstein-Barr virus-mediated B-cell immortalization. Proc Natl Acad Sci U S A 113, E782–790, doi:10.1073/pnas.1517141113 (2016).

